# Genomic insights into temperature-dependent transcriptional responses of *Kosmotoga olearia*, a deep-biosphere bacterium that can grow from 20°C to 79°C

**DOI:** 10.1101/060053

**Authors:** Stephen M. J. Pollo, Abigail A. Adebusuyi, Timothy J. Straub, Julia M. Foght, Olga Zhaxybayeva, Camilla L. Nesbø

## Abstract

Temperature is one of the defining parameters of an ecological niche. Most organisms thrive within a temperature range that rarely exceeds ∼ 30°C, but the deep subsurface bacterium *Kosmotoga olearia* can grow over a temperature range of 59°C (20°C -79°C). To identify genes correlated with this flexible phenotype, we compared transcriptomes of *K. olearia* cultures grown at its optimal 65°C to those at 30°C, 40°C, and 77°C. The temperature treatments affected expression of 573 of 2,224 *K. olearia* genes. Notably, this transcriptional response elicits re-modeling of the cellular membrane and changes in metabolism, with increased expression of genes involved in energy and carbohydrate metabolism at high temperatures and up-regulation of amino acid metabolism at lower temperatures. At sub-optimal temperatures, many transcriptional changes were similar to those observed in mesophilic bacteria at physiologically low temperatures, including up-regulation of typical cold stress genes and ribosomal proteins. Comparative genomic analysis of additional Thermotogae genomes, indicate that one of *K. olearia*'s strategies for low temperature growth is increased copy number of some typical cold response genes through duplication and/or lateral acquisition. At 77°C one third of the up-regulated genes are of hypothetical function, indicating that many features of high temperature growth are unknown.

## Introduction

Microorganisms are capable of growing over an impressive temperature range, at least from -15°C to 122°C (Takai et al. 2008;Mykytczuk et al. 2013), and temperature is one of the most important physical factors determining their distribution, diversity, and abundance (Schumann 2009). However, individual microbial species grow only within a much narrower temperature interval. For example, *Escherichia coli* O157:H7 thrives in the laboratory between 19°C and 41°C (Raghubeer and Matches 1990), while *Geobacillus thermoleovorans* has a growth range of 37°C to 70°C (Dinsdale et al. 2011). Microorganisms with temperature ranges >50°C are rare and, to date, research into the few that have ranges >40°C has focused on psychrophiles (e.g. Mykytczuk et al. 2013). *Kosmotoga olearia* TBF 19.5.1 (hereafter referred to as *K. olearia*) is an anaerobic thermophile from the bacterial phylum Thermotogae with a growth range that spans almost 60°C (DiPippo et al. 2009). How does this organism achieve such physiological flexibility, and what are the evolutionary advantages and implications of this capability?

Fluctuations in temperature induce broad physiological changes in cells, including growth rate, alterations to cell wall and membrane composition, translation, and energy metabolism (Barria et al. 2013;Pollo et al. 2015;Schumann 2009). These physiological changes can be classified into two broad types of cellular response. Cold or heat *shock* designates the changes observed *immediately* after the shift of a culture to a lower or higher temperature, while *prolonged growth* at a specific lower or higher temperature elicits an *acclimated* low- or high-temperature response (Barria et al. 2013;Schumann 2009). Most studies of prokaryotes have focused on temperature shock responses rather than acclimated growth. Among the Thermotogae, responses to both heat shock and prolonged growth at high temperatures have been studied in the hyperthermophile *Thermotoga maritima*, which can grow between 55°C and 90°C (Pysz et al. 2004;Wang et al. 2012). During prolonged high-temperature growth *T. maritima* strongly up-regulates central carbohydrate metabolism genes and expresses a few typical heat shock protein genes (Wang et al. 2012). Little is known about how *T. maritima* responds to sub-optimal temperatures, although it encodes some genes implicated in cold shock response. For example, its family of cold shock proteins (Csp), which are nucleic acid chaperones known to be induced during cold shock and cold acclimation in mesophilic bacteria (Barria et al. 2013;Phadtare 2004), exhibits nucleic acid melting activity at physiologically low temperatures (Phadtare et al. 2003). Similarly, responses to cold shock in a few other thermophiles involve many of the genes implicated in mesophilic cold shock response (e.g. Boonyaratanakornkit et al. 2005;Mega et al. 2010). In this study, we systematically assessed bacterial physiological changes associated with response to prolonged growth at both high and low temperature using *K. olearia* as a model system. Such changes can reflect not only response to temperature itself, but also growth rate effects and general responses to stress. By conflating these factors, our study examined overall changes in *K. olearia*'s gene expression in the environments defined by a specific temperature.

*K. olearia* was isolated from a deep subsurface oil reservoir with *in situ* temperature of 68°C (DiPippo et al. 2009). Cultivation experiments, 16S rRNA surveys, and metagenomic analyses have confirmed that the principal natural habitat of *K. olearia* is deep subsurface sediments and that these bacteria are particularly abundant in high temperature petroleum reservoirs (e.g. Vigneron et al. 2017;Duncan et al. 2009;Gittel et al. 2009;Pollo et al. 2016;Nesbø et al. 2010;Kotlar et al. 2011). However, *K. olearia* has also been either isolated or detected via 16S rRNA sequencing in oil fields (Nesbø et al. 2010), in an underground salt cavern plant used to store oily sand (Bordenave et al. 2013), and in engineered systems (Duncan et al. 2014;Oko et al. 2017) with *in situ* temperatures of 20°C–50°C, indicating that this bacterium occupies an ecological niche defined by a broad temperature range.

The *K. olearia* genome (NC_012785) has 2,302,126 bp and is predicted to encode 2,224 genes (Swithers et al. 2011). Within the Thermotogae, genome size, intergenic region size, and number of predicted coding regions correlate with the optimal growth temperature of an isolate (Zhaxybayeva et al. 2012), with hyperthermophilic Thermotogae genomes being the most compact. Phylogenetically, the Thermotogae order Kosmotogales comprises the genera *Kosmotoga* and *Mesotoga* spp., the latter being the only described mesophilic Thermotogae lineage (Pollo et al. 2015). Assuming a hyperthermophilic last common ancestor of the Thermotogae (Zhaxybayeva et al. 2009), the Kosmotogales can be hypothesized to have acquired wide growth temperature tolerance secondarily by expanding its gene repertoire. Moreover, it is likely that the ability of the Kosmotogales common ancestor to grow at low temperatures enabled the evolution of mesophily in *Mesotoga* (Pollo et al. 2015).

Such adaptations to novel environments can be greatly facilitated by lateral gene transfer (LGT), since genes already "adapted" to the new conditions are readily available in the microbial communities of the new environment (Boucher et al. 2003). For instance, LGT has been implicated in adaptation to high temperature growth in hyperthermophilic bacteria, including *Thermotoga* spp., and to low temperature growth in archaea (López-García et al. 2015;Pollo et al. 2015;Boucher et al. 2003). Genome analysis of the mesophilic *Mesotoga prima* revealed that it laterally acquired 32% of its genes after it diverged from other Thermotogae (Zhaxybayeva et al. 2012). Many of the predicted gene donors are mesophiles, supporting the importance of lateral acquisition of genes already adapted to mesophilic conditions in the evolution of *Mesotoga*.

To further gain insights into mechanisms of bacterial temperature response we sequenced 13 transcriptomes from isothermal cultures of *K. olearia* and examined transcriptional differences at temperatures spanning its wide growth range (30°C, 40°C, 65°C, and 77°C). We also investigated the importance of gene family expansion for adaptation of *K. olearia* to growth over a wide temperature range via comparative genomic and phylogenetic analyses of identified temperature responsive genes and their homologs in two newly sequenced *Kosmotoga* isolates, as well as in genomes of other thermophilic and mesophilic Thermotogae.

## Results

### Temperature shifts and isothermic conditions elicit different growth patterns in *K. olearia*

Under laboratory conditions in liquid anaerobic medium we observed growth of *K. olearia* at temperatures as low as 25°C and as high as 79°C, with optimal growth at 65°C, defined as the temperature affording the fastest growth rate (Fig. 1 and Fig. S1). Using a non-linear regression model (Ratkowsky et al. 1983) we estimate a growth-permissive temperature range of 20.2 – 79.3°C, consistent with the previously reported wide growth range of this isolate (DiPippo et al. 2009). Interestingly, we were not able to cultivate *K. olearia* at temperatures near its range boundaries (30°C and 77°C) by direct transfer from 65°C cultures. Instead, the growth temperature had to be changed sequentially in ≤10°C increments. Particularly at the extremes, even small temperature shifts caused both a longer lag phase and a slower growth rate compared to isothermal cultures (Fig. 1 and Fig. S1). This phenomenon has been previously noted for mesophilic bacteria, especially for transitions from high to low temperature (Swinnen et al. 2004). Our observations suggest that cells shifted to a new temperature need to undergo large physiological changes that require time (i.e. an ‘acclimation’ period (Barria et al. 2013)), and that these physiological challenges are too great to overcome when temperature changes are large.

**Fig. 1.**
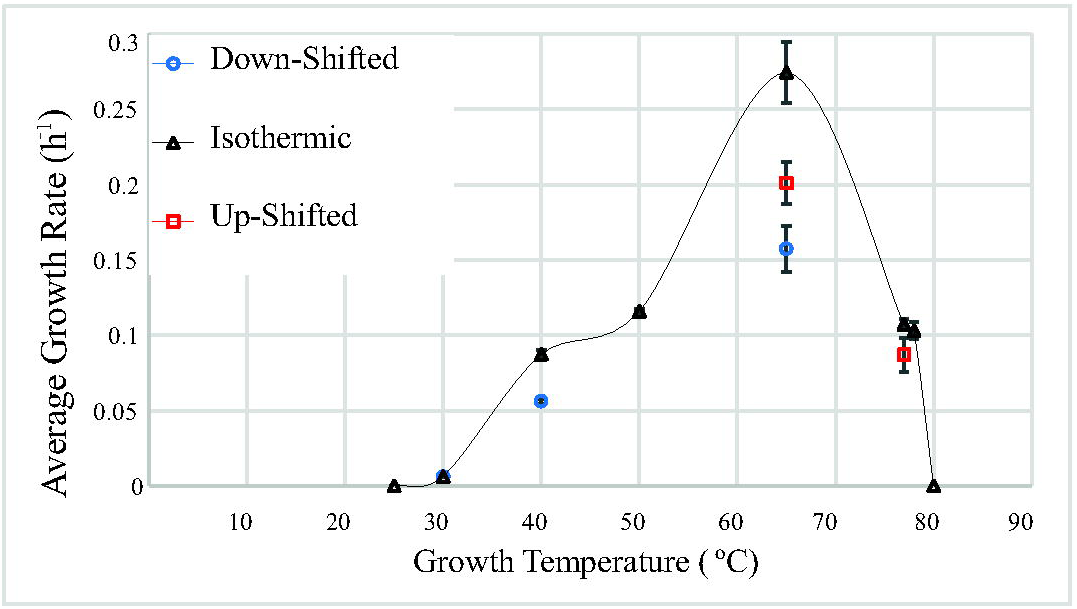
Growth rate of *K. olearia* as a function of temperature. Isothermic growth curves were generated at each temperature from an inoculum grown at that temperature for at least three transfers (except for 25°C and 80°C, for which an inoculum from the same temperature could not be generated; see materials and methods). Up-shifted and down-shifted growth curves were generated from an inoculum that was grown at lower and higher temperatures, respectively. Red squares, growth temperature up-shifted from 65°C to 77°C or from 40°C to 65°C; Blue circles, growth temperature down-shifted from 77°C to 65°C, 65°C to 40°C, or 40°C to 30°C. Data points represent the mean of replicate cultures (see materials and methods); error bars represent standard error.

### Architecture of the *K. olearia* transcriptome

Analysis of transcription start and stop sites predicted a minimum of 916 transcriptional units (TU) in *K. olearia* (Supplemental material and Table S2), 52% of which consist of a single gene. This fraction of single-gene TUs lies between the 65% reported for *E. coli* (Cho et al. 2009) and the 43% recorded for *T. maritima*, which has also been shown to have a streamlined genome and a low-complexity transcriptome (i.e. few sub-operonic transcripts and few genes with multiple start sites) (Latif et al. 2013). The average TU length of ∼2.39 genes in *K. olearia* is less than the 3.3 genes per transcript of *T. maritima* (Latif et al. 2013) but closer to 2.2 genes per transcript in the mesophilic firmicute *Geobacter sulfurreducens* (Qiu et al. 2010) and 1-2 genes per transcript in bacteria in general (e.g. Cho et al. 2009)). Given that the *K. olearia* genome has more intergenic DNA than *T. maritima*'s genome (the ratio of the nucleotides located in non-coding vs. coding regions is 0.13 in *K. olearia* and 0.06 in *T. maritima*), the shorter TU lengths in *K. olearia* may point to more flexible transcriptional regulation.

### Consistent energy generation across different temperature conditions

*K. olearia* produces ATP from pyruvate using a biochemically well-understood fermentation pathway that generates hydrogen, carbon dioxide and acetate ((DiPippo et al. 2009); Fig. 2 and data not shown). Since pyruvate was the carbon and energy source provided in all experiments, we surveyed 51 genes predicted to be involved in core energy metabolism during growth on pyruvate. Based on expression levels, we constructed a model that accounts for all major end products during growth at 65°C (Fig. 2) and contains 15 genes with consistently high expression across all temperature treatments (Fig. S2a and Table S3). In addition to indirectly validating the previously known functional annotations of these genes, we furthermore propose that the most highly expressed ABC-transporter gene cluster, Kole_1509 – 1513, encodes a pyruvate transporter (Fig. 2). Its current annotation as a peptide ABC transporter may be erroneous, since most of the peptide ABC transporters predicted in *T. maritima* using bioinformatics have been shown instead to bind and transport sugars (Nanavati et al. 2006). Further functional studies of the transporter (e.g. binding assays, expression with different substrates) are needed to confirm this hypothesis.

**Fig. 2.**
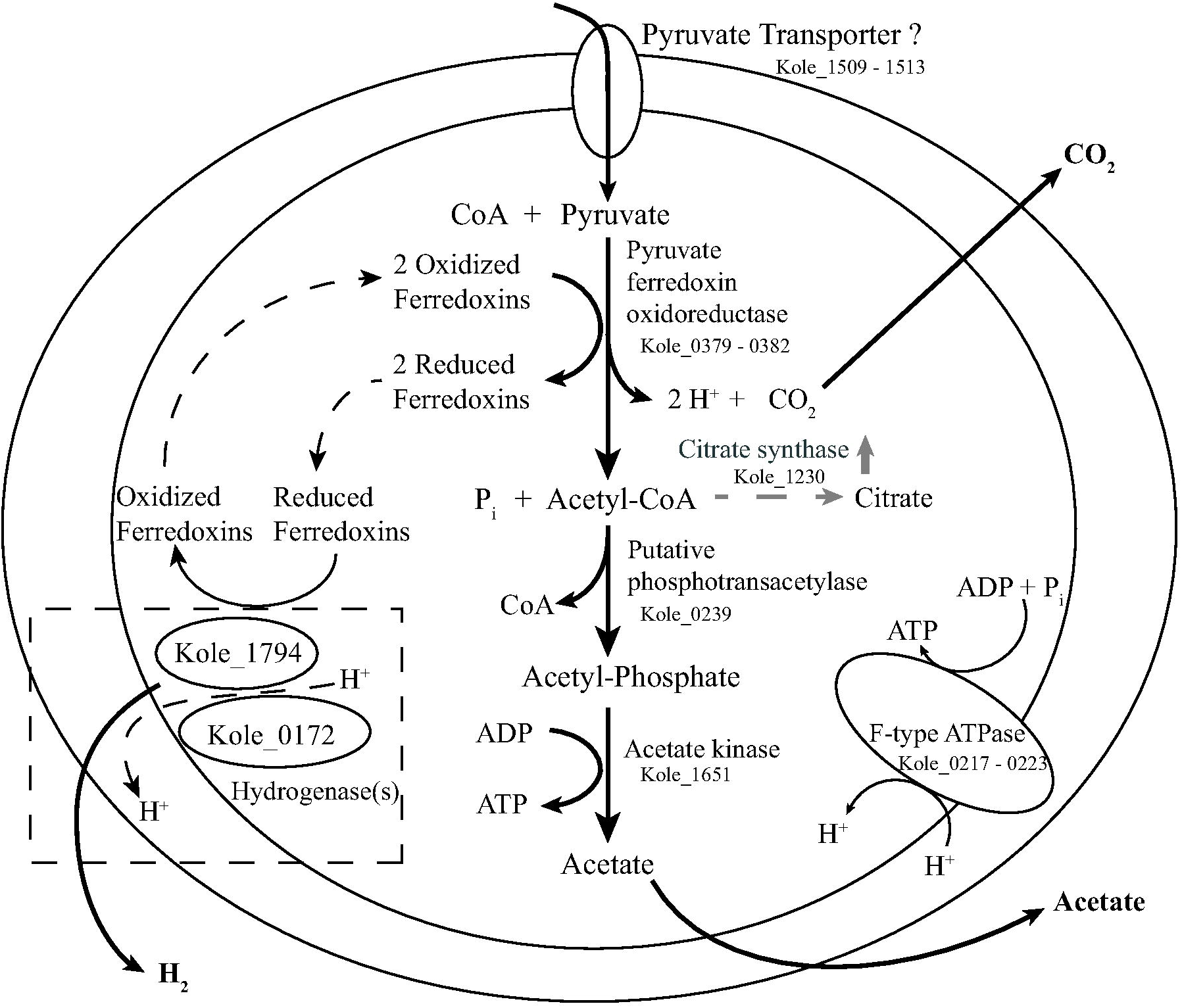
Model of energy generation pathway in *K. olearia* during growth on pyruvate. The model includes genes likely involved in conversion of pyruvate to acetate, CO_2_, H_2_, and ATP. The genes were selected from the list of genes highly expressed across all temperature conditions (Fig. S2a, Table S3). Acetate transport is not shown. The dashed box indicates hydrogenase activity. The two highly expressed hydrogenases are shown, but their potential interactions with each other or with the membrane are not known. Increased expression of citrate synthase at low temperature, which could redirect acetyl-CoA away from acetate production, is shown in grey. The assignment of the ABC transporter as pyruvate transporter (Kole_1509-1513) is based on its high expression level, but experiments are needed to confirm its involvement in pyruvate transport. The model also explains the observed lower ratio of carbon dioxide to hydrogen produced by growth on maltose vs. pyruvate (not shown), because during growth on maltose reduced electron carriers would be generated from the conversions of maltose to pyruvate.

### Identification of temperature-related transcriptional responses in *K. olearia*

Hierarchical clustering and Principal Component Analysis (PCA) separated the transcriptomes according to temperature treatment (Fig. 3, Fig. S3 and Supplemental material), suggesting that the observed changes in transcription are due to the culture growth temperature. Several genes with a high correlation between their expression level and a specific growth temperature (Table S4 and vectors in Fig. 3) are known to be involved in temperature response (Pollo et al. 2015). For example, expression of the heat shock serine protease Kole_1599 positively correlated with the 77°C transcriptomes (Fig. 3), where high expression of this protease was expected based on its involvement in heat shock response in *T. maritima* (Pysz et al. 2004). Similarly, expression of the cold shock protein genes Kole_0109 and Kole_2064 positively correlated with low temperature growth (Fig. 3). Lastly, some observed changes presumably were due to the expected decreased metabolic activity of the culture at sub- and supra-optimal temperatures. This can be exemplified by the high expression and strong correlation of the alcohol dehydrogenase Kole_0742 (see Supplemental material for further discussion of the expression pattern of this gene) and the central carbon metabolism gene glyceraldehyde-3-phosphate dehydrogenase (Kole_2020) with the 65°C transcriptomes.

**Fig. 3.**
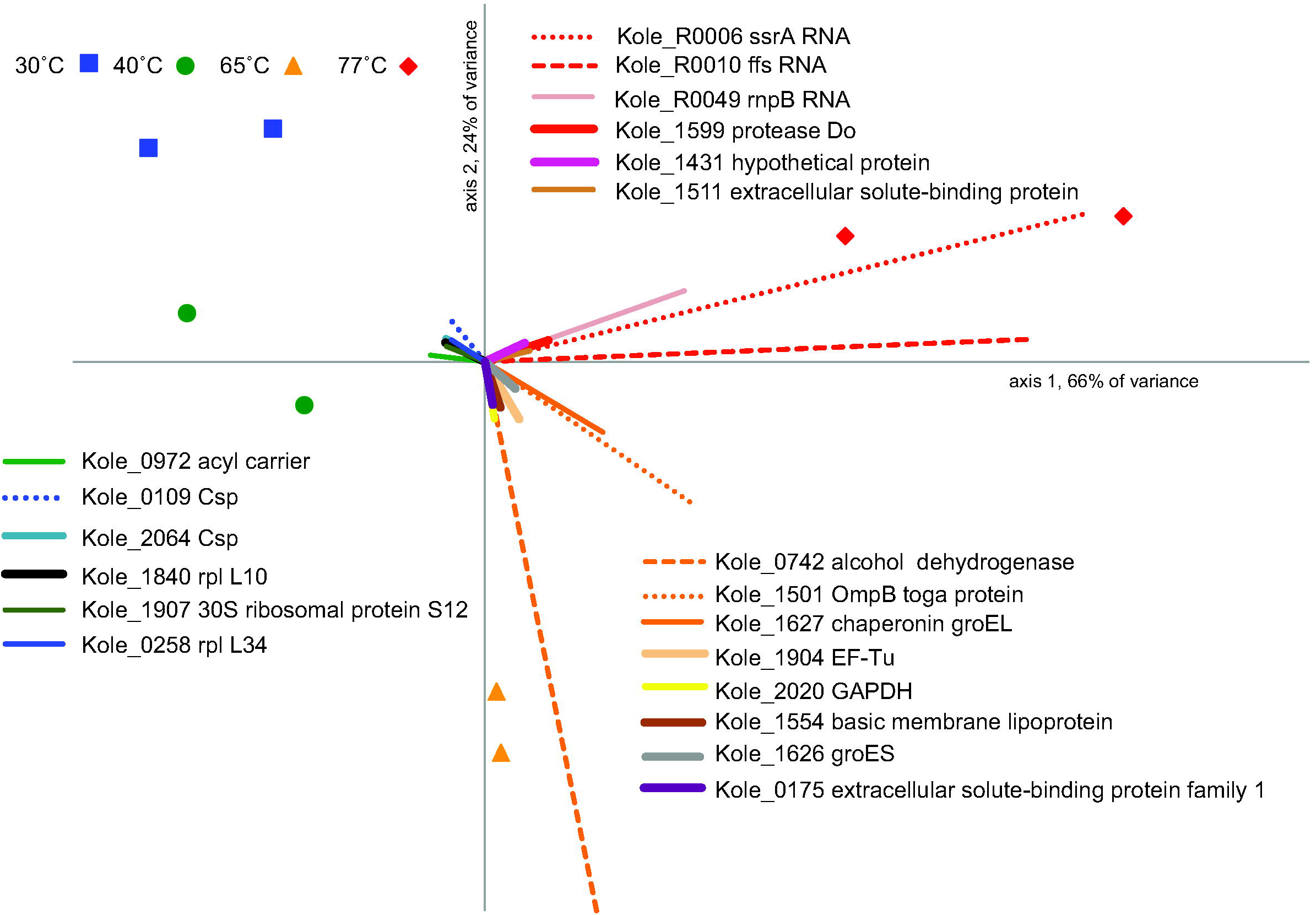
Biplot of the principal component analysis of 8 transcriptomes. The plot is based on expression values from all genes. Each transcriptome is denoted by a point and genes are represented by vectors. The 20 longest (i.e. most highly correlated) gene vectors are shown. Coordinates and vector length for all genes can be found in Table S4. It should be noted that the *ffs* (Kole_R0010) transcript is only 115 nt, and may not have been fully represented in every transcriptome due to our isolation protocol which selects against small RNA (<200 nucleotides). Also, the high expression of the alcohol dehydrogenase (Kole_0742) is probably due to the addition of stop solution before RNA isolation (see Supplemental material).

Putative temperature-responsive genes were identified by pairwise comparisons of Illumina transcriptomes from each temperature treatment to the optimal condition of 65°C (i.e. 30°C vs 65°C, 40°C vs 65°C, and 77°C vs 65°C). Across all comparisons 573 genes fulfilled our criteria for temperature responsiveness (≥ 2-fold difference in expression, > 20 reads per transcript, False Discovery Rate < 0.05) with 430, 115, and 169 genes detected in the 30°C vs 65°C, 40°C vs 65°C, and 77°C vs 65°C comparisons, respectively (Table S5). Expression of 306 of the 573 putative temperature-responsive genes correlated with growth rate (r > |0.7|; Table S5). We used ANCOVA to untangle the effect of growth rate from temperature, which revealed that growth rate was a significant factor for only 60 of the 306 genes (p < 0.05; Table S5). Moreover, for 26 of these 60 genes temperature was also a significant factor, and the simpler growth-rate-only model was rejected for all but 13 of the 60 genes (Likelihood Ratio Test, FDR-corrected p-value < 0.05; Table S5). In an alternative comparison based on relative effect size of temperature vs. growth rate, growth rate had larger influence on expression of 51 of the 306 genes (Table S5). Among these genes were the glyceraldehyde-3-phosphate dehydrogenase (Kole_2020) and the iron-containing alcohol dehydrogenase (Kole_0742) mentioned above. Therefore, although we retained the designation of "putatively temperature-responsive" for all 573 genes, below we focus on genes not primarily affected by growth rate.

### Distribution of temperature responsive genes across functional categories suggests a regulated response to temperature

Most of the temperature-responsive genes were up-regulated when compared to the expression at 65°C (155 genes are down-regulated and 559 up-regulated; Table S5). One notable exception to this overall trend were genes involved in carbohydrate and energy metabolism (Clusters of Orthologous Groups [COG] categories C and G) where 32 genes were down-regulated at 30°C compared to 15 genes up-regulated. In transcriptomes at all non-optimal temperatures the list of putative temperature-responsive genes was depleted in genes involved in translation (COG category J) and nucleotide metabolism (COG category F) (Fig. 4) and conversely enriched in genes involved in signal transduction (COG category T), and replication, recombination and repair (COG category L, particularly at 30°C). Differential expression of the signal transduction genes (COG category T) suggests the importance of these systems for regulating cellular responses at all tested temperatures. Most of the identified COG category L genes are either mobile elements or CRISPR-associated proteins, hinting at an increased activity of mobile genetic elements – a known common feature of stress responses (Foster 2007). Additionally, at both 30°C and 77°C many genes encoding transcription regulators (COG category K) are up-regulated, implying that prolonged growth at sub- and supra-optimal temperatures results in detectable changes in transcriptional gene regulation in *K. olearia*.

**Fig. 4.**
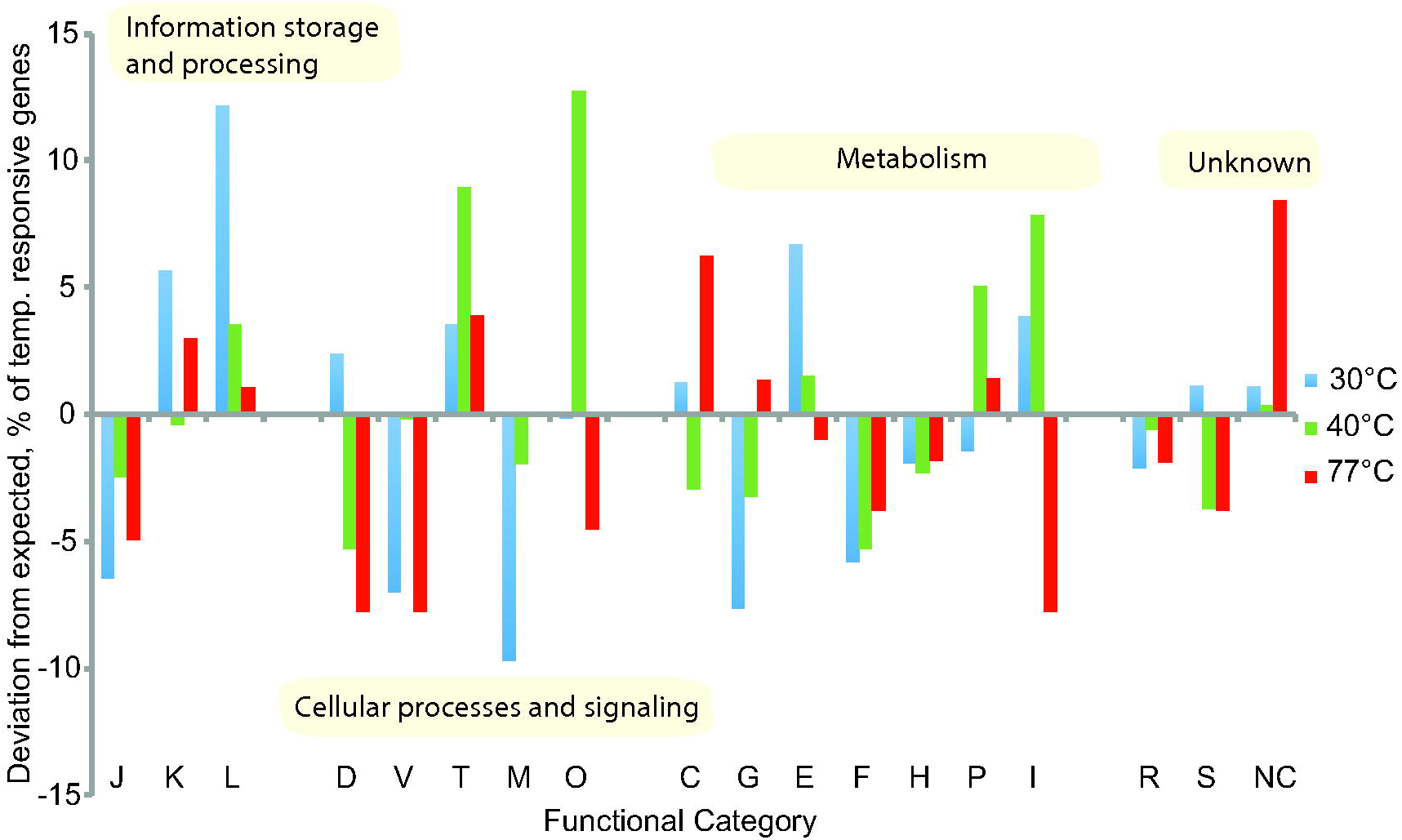
Difference between observed and expected number of temperature responsive genes across functional categories. Functional categories were assigned using the Clusters of Orthologous Groups (COG) database as implemented in IMG (Markowitz et al. 2014) and are denoted by one-letter abbreviations along the X-axis. For each temperature treatment (30°C, 40°C and 77°C) only the temperature-responsive fraction of the *K. olearia* genome was considered. If the temperature-responsive genes were randomly distributed across functional categories we would expect the same fraction of temperature-responsive genes in each COG category. The difference (in percent) between the observed and expected number of temperature responsive genes is plotted on the Y-axis with positive and negative values referring to over- and under-representation of the temperature-responsive genes, respectively. COG category abbreviations: J: Translation, ribosomal structure and biogenesis, K: Transcription, L: Replication, recombination and repair, D: Cell cycle control, cell division, chromosome partitioning, V: Defense mechanisms, T: Signal transduction mechanisms, M: Cell wall/membrane/envelope biogenesis, U: Intracellular trafficking, secretion, and vesicular transport, O: Post-translational modification, protein turnover, and chaperones, C: Energy production and conversion, G: Carbohydrate transport and metabolism, E: Amino acid transport and metabolism, F: Nucleotide transport and metabolism, H: Coenzyme transport and metabolism, I: Lipid transport and metabolism, P: Inorganic ion transport and metabolism, Q: Secondary metabolites biosynthesis, transport, and catabolism, R: General function prediction only, S: Function unknown, NC: Not in COG database.

### At 40°C there are pronounced differences in membrane fatty acid composition but no signs of cold stress

Although the growth rate of *K. olearia* at 40°C is only one-third of that at the optimum 65°C (Fig. 1 and Fig. S1), clustering analysis suggested that the 40°C transcriptome was most similar to that at 65°C (Fig. 3 and Fig. S3). However, at 40°C even the four most highly expressed temperature-responsive genes, including the growth-rate dependent alcohol dehydrogenase (Kole_0742), had significantly lower expression levels (Table S5), reflecting the slower metabolic rate at lower temperature growth. Yet, 94 of 115 putative temperature responsive genes were up-regulated (Table S5, Fig. S2b), suggesting that slower metabolism is not the only explanation for the observed transcriptional response to growth at 40°C.

Lipid metabolism (COG category I) appears to be particularly important at 40°C. For instance, all the predicted fatty acid synthesis genes showed the highest expression at 40°C (Fig. S4), with significantly higher expression of two genes involved in synthesis of unsaturated fatty acids (Kole_0968) and initiation of fatty acid synthesis (Kole_0969). Biochemical analyses of total fatty acids at 40°C and 65°C showed a much greater diversity of fatty acids at 40°C (Table S6), which may explain the higher demand for the products of these genes at lower temperatures. Interestingly, there was increased expression of a phosphate ABC transporter (Kole_0707 – Kole_0711, Table S5, Fig. S2b), which may be linked to increased production of polar membrane lipids at moderately low temperatures.

An enrichment of differentially expressed genes in “post-translational modification, protein turnover, and chaperone function” (COG category O; the category that harbors genes related to cellular stress) was due to both up- and down-regulation of the involved genes (Fig. 4). For instance, *K. olearia* has three temperature-responsive peptidylprolyl isomerase (PPIase) genes: two PpiC-type genes (Kole_1682 and Kole_0383) that are both highly expressed at 40°C, and one FKBP-type gene (Kole_1745) that shows high expression at all temperatures except 77°C (Table S5). At lower temperatures (e.g. 37°C), these enzymes catalyze proline isomerization, which happens spontaneously at higher temperatures (Godin-Roulling et al. 2015). However, the enzymes known to assist protein folding under cellular stress, chaperones (GroEL [Kole_1627] and Hsp70 [Kole_0886]) and protease Do (Kole_1599), were significantly down-regulated at 40°C (Table S5 and Fig. S2b). Among other typical cold stress-related proteins, only one of *K. olearia*’s three cold shock proteins (Kole_0109) showed significantly higher expression at 40°C, but its up-regulation was merely moderate when compared to its expression levels at 30°C (Table S5 and Fig. S2b). Taken together, the observed expression patterns of known cold stress-related genes are consistent with the cells being in a non-stressed state at 40°C.

### *K. olearia* is in cold stress at 30°C

Among the three most highly expressed up-regulated genes at 30°C are two Csp-encoding genes (Kole_0109 and Kole_2064, Table S5, Fig. S2c), suggesting that the cells were in a cold-stressed state during growth at 30°C. In support of this there was also significant up-regulation of other genes previously linked to bacterial cold response (e.g. Alreshidi et al. 2015;Barria et al. 2013), including a DEAD/DEAH-box RNA helicase (Kole_0922), *rbfA* (Kole_2103), *nusA* (Kole_1529) and ribosomal proteins L10 (Kole_1840) and L7/L12 (Kole_1839). Genes encoding several additional ribosomal proteins and ribosomal RNA (rRNA) methyltransferases, *rmlH* (Kole_1718) and *rmlL* (Kole_0897), were also up-regulated (Fig. 3 and Table S5).

To detect a decrease in environmental temperature and elicit an appropriate regulatory response, some bacteria have evolved two-component cold sensors (de Mendoza 2014). These signal transduction systems consist of a sensor, a membrane-integrated protein with a kinase domain that detects changes in the fluidity of the cell membrane, and the cytoplasmic response regulator, a protein that induces expression of cold-responsive genes. In *K. olearia*, a histidine kinase with two predicted transmembrane domains (Kole_1017) and two response regulators (Kole_1015 and Kole_1016) showed a steady increase in expression as temperatures decreased from 65°C, but no significant change in expression at 77°C (Table S5), leading us to hypothesize that these genes encode a cold-sensing two-component system.

At 30°C, and to a lesser extent at 40°C, we also observed an over-representation of highly expressed genes involved in amino acid metabolism (COG category E). Specifically, several genes in the arginine (Kole_0092 – Kole_0097) and lysine (Kole_0104 – Kole_0107, 30°C only) biosynthesis pathways were up-regulated, suggesting the potential for accumulation of peptides and amino acids (or their intermediates) at lower temperatures. Interestingly, while the cells may accumulate peptides at 30°C, at 40°C there was increased expression of an oligo-peptidase (Kole_1190) and genes involved in lysine degradation (Kole_0958, Kole_0963 – Kole_0966). Such distinguishably different metabolic responses to moderately low (40°C) and low (30°C) temperatures suggest a fine-tuned temperature-dependent peptide turnover.

### *K. olearia* is in heat stress at 77°C

Since 77°C is near the upper limit for *K. olearia* growth under our laboratory conditions, we hypothesize that the observed differences in expression profiles at 65 and 77°C would reflect a cell-wide heat stress response. Of the 169 significantly differentially expressed genes, 119 showed increased expression at 77°C (Table S5 and Fig. S2d). Among the most highly expressed genes were those encoding the structural RNAs *ffs* (Kole_R0010), *ssrA* (Kole_R0006), and *rnpB* (Kole_R0049) (Fig. 3), suggesting an increased rate of RNA turnover at supra-optimal temperature. Moreover, genes involved in carbohydrate and energy metabolism (COG C and G categories) were over-represented and up-regulated at 77°C (Fig. 4, Table S5). However, only two of the known heat *shock* response genes (Pysz et al. 2004), the extreme heat stress sigma factor-24 (*rpoE*, Kole_2150) and the heat shock protease (Kole_1599), were up-regulated, and 41 of 119 genes (34%) are annotated as “hypothetical proteins”, indicating that adaptation to growth at sustained high temperature remains largely uncharacterized. Putative functions could be inferred for some of the encoded proteins of these 41 genes of unknown function. Seventeen of the 41 proteins have predicted signal peptides, suggesting that they may be secreted, and four have two or more predicted transmembrane regions, suggesting they are membrane-associated. Kole_0652 carries a PrcB-domain, which interacts with and stabilizes PrtP protease (Godovikova et al. 2010). Kole_1314 (and its paralog Kole_1297) contains an AbiEii-toxin domain, and may be part of a toxin-antitoxin system. Furthermore, the 41 genes could be grouped into 34 transcriptional units, each containing at most three of these genes (Table S2) and scattered across the *K. olearia* genome. Two of the genes, Kole_1430 and Kole_1431, are co-transcribed with genes from a two-component system (Kole_1428 and Kole_1429), suggesting they may be involved in environmental sensing or signaling. Three other co-transcribed genes (Kole_1266, Kole_1267, Kole_1270) are found in a cluster containing CRISPR-genes, and two of them (Kole_1266 and Kole_1270) contain RAMP-domains, suggesting CRISPR-related function (Makarova et al. 2011). The majority of the 41 genes of unknown function have homologs in genomes of other *Kosmotoga* spp. (N=38), *Mesotoga* spp. (N=23), and other Thermotogae (N=26).

### Global regulators of temperature response

The transcriptional changes seen at the sub- and supra-optimal temperatures are likely to be controlled by one or a few global regulators (Balleza et al. 2009), such as some temperature sensors (de Mendoza 2014) and sigma factors needed for transcription initiation (Buck et al. 2000). The two-component cold sensor up-regulated at low temperatures (Kole_1015 - Kole_1017) may represent one such global regulator. Two different sigma factors (Kole_2150 and (Kole_1408) were significantly up-regulated at 77°C and at 30°C and 40°C, respectively (Table S5), hinting at existence of temperature-specific sigma factors. Kole_2150 belongs to the sigma-24 ECF subfamily, which is activated in response to environmental stress (Balleza et al. 2009). Kole_1408 belongs to the sigma-54 family, which is involved in enhancer-dependent transcription (Buck et al. 2000), introducing the possibility that this sigma factor may be a target of the two-component cold sensor. In general, at both 30°C and 77°C the differentially expressed genes were enriched in genes involved in transcriptional regulation (COG category K) (Fig. 4), leaving a possibility of additional global regulators.

### Conservation of *K. olearia*’s temperature-responsive genes within the *Kosmotoga* genus

All genes that are required for adaptation and response of *K. olearia* strain TBF 19.5.1 to a wide range of growth temperatures are expected to be present in other *K. olearia* isolates, whereas some may be absent from *Kosmotoga* species that have a narrower spectrum of growth temperature. Therefore, we contrasted the *K. olearia* genome to the genomes of *Kosmotoga* sp. DU53 and *Kosmotoga arenicorallina* (Pollo et al. 2016). When compared to *K. olearia*, *Kosmotoga* sp. DU53 has a similar growth temperature range (25°C - 79°C, Table S7) and >99% average nucleotide identity (ANI), while *K. arenicorallina* exhibits a narrower growth temperature range (35°C - 70°C, Table S7) and has only 84% ANI.

Indeed, the *Kosmotoga* sp. DU53 genome lacks only 10 of the 573 *K. olearia* putative temperature-responsive genes (BLASTP and TBLASTN searches, E-value <10^−3^, Table S5). All 10 genes were expressed in *K. olearia* at relatively low levels, suggesting that they are unlikely to be essential for high or low temperature growth. On the other hand, the *K. arenicorallina* genome does not have detectable homologs of 103 of the 573 *K. olearia*’s putative temperature-responsive genes (BLASTP and TBLASTN searches, E-value <10^−3^; Table S5). The list of absent genes includes several of the arginine and lysine biosynthesis genes that are up-regulated in *K. olearia* during growth at 30°C, and seven of the genes of unknown function up-regulated at 77°C. Therefore, we hypothesize that a subset of these 103 genes may play a role in extending the growth range of *K. olearia* to ≤35°C and ≥70°C.

### Many *key* temperature-responsive genes are laterally acquired

Obtaining "pre-adapted" genes from other genomes is one way prokaryotes adjust to new environmental conditions (Boucher et al. 2003). Using HGTector (Zhu et al. 2014) we predicted that 354 of *K. olearia*’s 2,118 protein coding genes have been acquired laterally by *K. olearia* or the Kosmotogales (i.e. *Kosmotoga* and *Mesotoga* genera), presumably representing LGT events occurring after the divergence of Kosmotogales from other Thermotogae (Table S8). Eighty-eight of the 354 genes were temperature responsive (Table S5, Fig. S5A and S5B), including several above-discussed highly expressed genes (Table 1), and 37 of the 88 genes are shared with *Mesotoga* (Table S5). Notably, 76% of these 37 genes are upregulated at 30°C (Fig. S5C), suggesting that their acquisition may have been important in adaptation to low temperature growth. Among these are the earlier mentioned rRNA methyltransferase genes (Kole_1718 and Kole_0897). The fatty acid synthesis genes (Kole_0969 - Kole_0973) that are up-regulated at 40°C, as well as their Kosmotogales homologs, form a distantly related sister clade to other Thermotogae lineages (Fig. S6A), implying that these genes may have been acquired from an un-sampled lineage. Similarly, the Csp-encoding gene highly expressed at 30°C (Kole_0109) is placed outside of the Thermotogae clade (Fig. S6B).

**Table 1.**
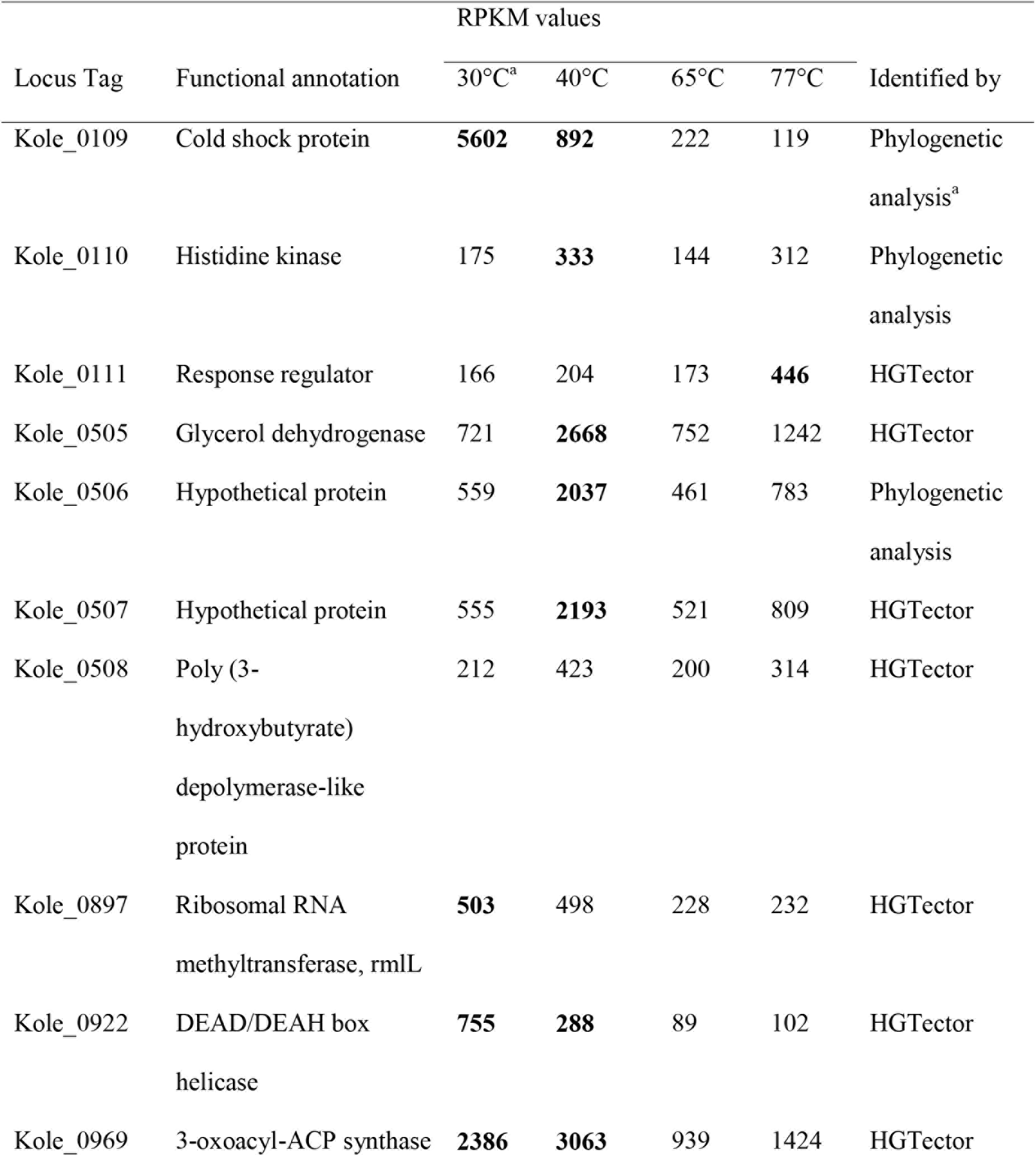
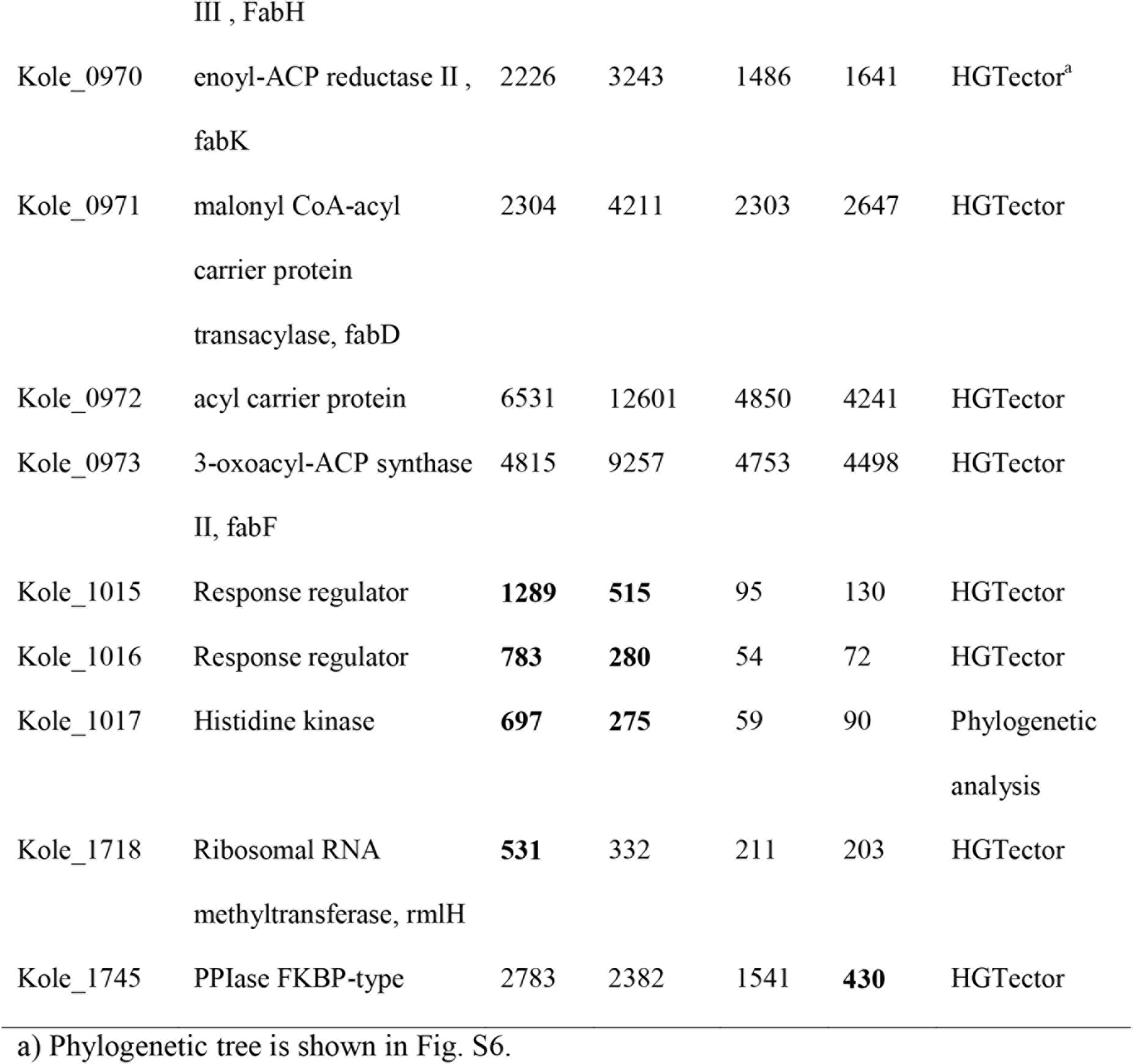
Gene expression in selected laterally acquired temperature-responsive genes. At each temperature, the listed RPKM values represent the average expression levels across replicates. Values that are significantly different from 65°C are shown in bold font. None of the temperature responsive expression patterns show significant effect of growth rate and, where tested (i.e. |R| > 0.7), the growth-rate-only model was rejected.

Notably, some putative lateral gene acquisitions by *K. olearia* do not have homologs in *Mesotoga*. These include genes encoding the predicted cold temperature sensor (Kole_1015 – Kole_1017), one of the PPIase genes (Kole_1745), as well as the canonical cold response enzyme DEAD/DEAH box RNA helicase (Kole_0922). Absence of these genes in *Mesotoga* suggests their potential importance for *K. olearia*'s ability to grow over a wide temperature range.

## Discussion

### How can *K. olearia* grow at such a wide range of temperature?

Examination of *K. olearia*'s transcriptional response to sustained exposure to a non-optimal temperature revealed both high expression of core metabolic genes and differential expression of hundreds of other genes, with selected features highlighted in Fig. 5. At each tested temperature, core metabolism genes for pyruvate utilization show high relative expression, which strongly suggests that *K. olearia* uses the same proteins for its core energy metabolism and that these proteins can function across its wide growth temperature range. In contrast, genes involved in regulatory functions showed significant changes in expression at all temperatures, particularly at 30°C and 77°C (COG category K and T in Fig. 4), suggesting that regulating gene expression is important in response to a temperature shift. Among these genes we identified putative global temperature regulators: the two-component cold sensor and temperature-specific sigma factors.

**Fig. 5.**
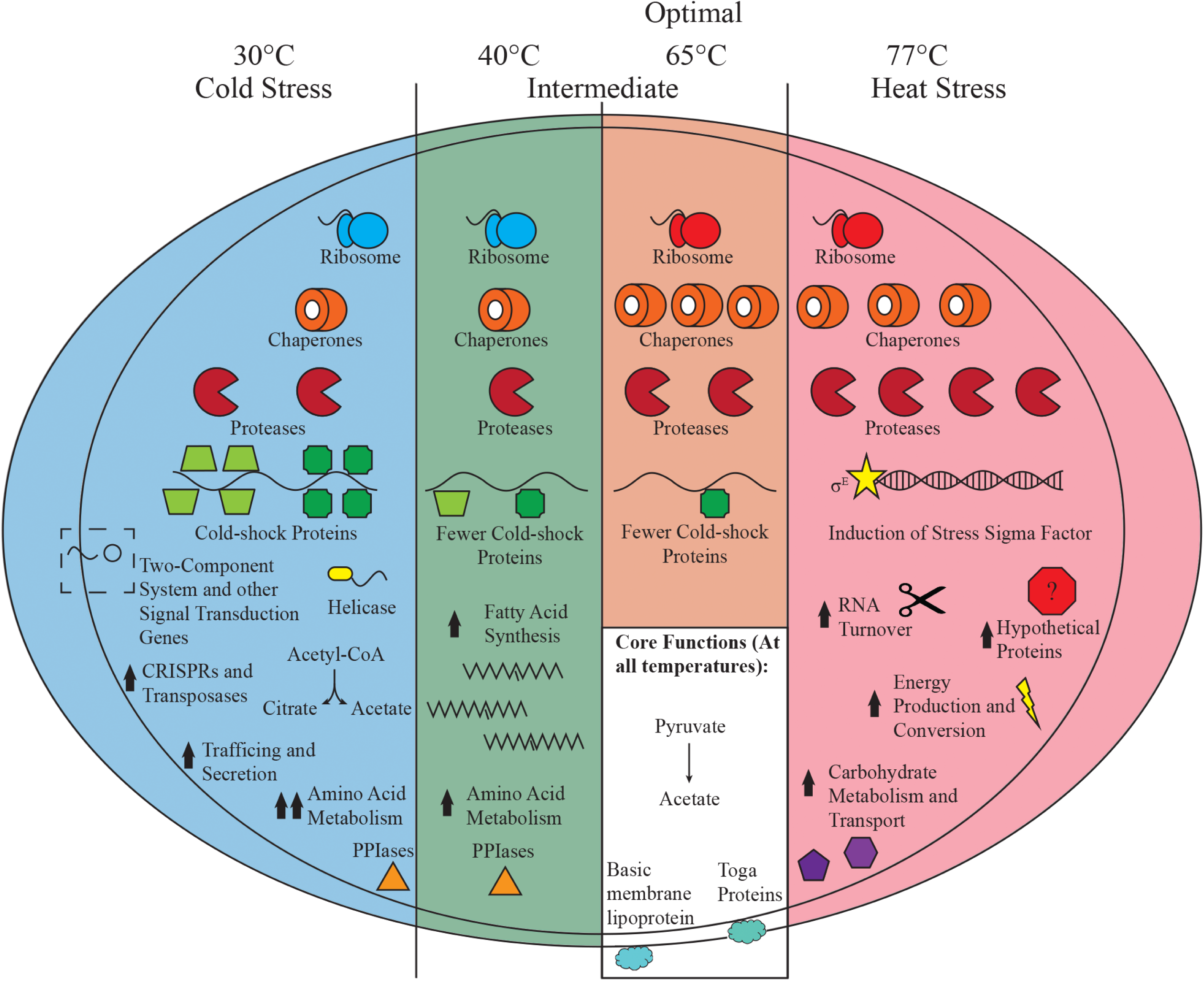
Schematic of *K. olearia*'s major temperature-induced transcriptional responses. Major responses outlined in the text occurring in the three states observed are shown. The number of chaperone, protease, and cold-shock symbols reflects their relative expression at each temperature. Chaperones include groEL (Kole_1627), groES (Kole_1626), and dnaK (Kole_0886). Proteases refer to protease Do (Kole_1599) and protease La (Kole_0536). Cold-shock genes consist of Kole_2064 (dark green squares) and Kole_0109 (green trapezoids). The different coloured ribosomes represent changes in ribosomal protein composition at sub-optimal temperatures (see text for discussion). The putative two-component regulatory system (Kole_1015 – 1017) that had a dramatic increase in expression under cold stress is shown. Prominent changes in functional categories (COGs) for each temperature condition are also shown. Arrows indicate relative gene expression when compared to growth at optimal temperature. Core functions occurring at all temperatures are shown in the white box. Basic membrane lipoprotein and the major toga anchor protein refer to Kole_1554 and Kole_1500, respectively. See Fig. 2 for genes involved in pyruvate metabolism.

Close to *K. olearia*'s growth temperature maximum of 77°C, carbohydrate and energy metabolism genes (COG categories C and G) were up-regulated (Fig. 4). It is unclear, however, if the underlying cause is the increased turnover of enzymes at elevated temperatures, a demand for more enzymes due to increased carbohydrate catabolism, or a combination of these factors. Increased carbohydrate metabolism in response to prolonged growth at supra-optimal temperature has been observed previously in *T. maritima* (Wang et al. 2012), and therefore may be a common adaptation to high temperature growth in the Thermotogae. As observed for *K. olearia,* the prolonged supra-optimal temperature growth of *T. maritima* also did not involve up-regulation of typical heat-shock response proteins (Wang et al. 2012). This highlights the difference between cellular response to an immediate heat-shock and to prolonged growth at supra-optimal temperature (Balleza et al. 2009), and in general justifies classifying the cellular response to temperature into these two distinct categories.

At the moderately sub-optimal growth temperature of 40°C, *K. olearia* cells face physiological challenges of proper protein folding (Godin-Roulling et al. 2015), and of maintenance of a functional cell membrane (de Mendoza 2014). Our observation that at 40°C, despite the lower growth rate, lipid metabolism genes were among the most highly expressed genes suggests that changes to the cell membrane composition are one of the most important adaptations for survival of *K. olearia* at lower temperatures. Proper protein folding may require enzymatic assistance (Godin-Roulling et al. 2015), which may be achieved via *K. olearia*'s three PPIases. The significant up-regulation of the PPIase genes at 40°C suggest that they are particularly important at moderately low temperatures where the cells are still relatively active. However, the overall lack of induction of typical stress-related genes at 40°C, especially when compared to 30°C and 77°C, suggests that 40°C is within the "Goldilocks" temperature range for *K. olearia.*

At 30°C *K. olearia* is clearly under cold stress (Fig. 5), as evidenced by expression of genes known to be implicated in cold response. One of the strategies for maintenance of proper cellular function at non-optimal temperatures (Pollo et al. 2015) is accumulation of compatible solutes. Specifically, re-modelling of amino acid metabolism and possible accumulation of amino acids as compatible solutes at low temperatures has been observed in bacteria (e.g. (Dahlsten et al. 2014;Ghobakhlou et al. 2015)). The up-regulation of many genes involved in amino acid metabolism suggests that *K. olearia* may also accumulate amino acids or their intermediates for this purpose, especially at 30°C. At 30°C there was also significant up-regulation of a citrate synthase gene (Kole_1230), suggesting that *K. olearia* cells may accumulate citrate, as was observed in *Staphylococcus aureus* during prolonged cold stress (Alreshidi et al. 2015). Alternatively, citrate synthase, together with isocitrate dehydrogenase (Kole_1227), may be involved in converting pyruvate or acetyl-CoA to 2-oxoglutarate, a precursor for several amino acids including arginine, which has been suggested to accumulate in for instance *Clostridium* (Dahlsten et al. 2014). Interestingly, the genome of strictly mesophilic *M. prima* encodes more genes involved in amino acid metabolism than the genomes of *K. olearia* and other thermophilic Thermotogae (Zhaxybayeva et al. 2012). This supports the notion that the shift towards more amino acid metabolism in *K. olearia* may be an adaptation to low temperature growth and suggests that the mesophilic *Mesotoga* have taken this a step further by also *acquiring* more amino acid metabolism genes. Moreover, amino acid metabolism genes are among the most numerous bacterial genes laterally acquired by mesophilic archaea, which is hypothesized to reflect archaeal adaptation to low temperature growth (López-García et al. 2015).

Ribosomes also need to be fine-tuned (i.e. to adjust the stoichiometric proportions of accessory proteins) to function properly at low temperature (Barria et al. 2013). Consistently, we observed a change in expression of several genes encoding ribosomal proteins. The most dramatic differential expression, however, was observed for a ribosomal protein gene not yet connected to cold response (L34; Kole_0258, Fig. 3, Table S5). L34, a bacteria-specific ribosomal protein hypothesized to be a relatively recent addition to the evolving ribosome (Fox 2010), is required for proper ribosome formation (Akanuma et al. 2014). A *Bacillus subtilis* mutant lacking the L34 gene showed particularly slow growth at low temperature (Akanuma et al. 2012), suggesting a role for L34 in this condition. We also observed significant increased expression of rRNA methyltransferases, and methylation of rRNAs has been associated with responses to environmental stress, including temperature (Baldridge and Contreras 2014). Therefore, we hypothesize that *K. olearia* modifies its ribosome by changing stoichiometry of the ribosome components and by methylating rRNA. Time required for such ribosomal adjustments could also explain the longer lag phase following temperature shifts (Fig. S1).

Expansion of cold-responsive gene families may represent a common strategy for low temperature adaptation, as has been noted in many bacteria, especially in psychrophiles (e.g. (Piette et al. 2010). *K. olearia* exhibits the same trend. For instance, when compared to other Thermotogae, all three analyzed *Kosmotoga* genomes harboured more copies of Csp-encoding genes (Table S9). The observed gene family expansions, however, might be important not necessarily for low temperature growth alone, but instead for growth over a wide temperature interval. *Mesotoga* functions with only a single *csp* gene, demonstrating that having more copies of this gene is not required for low temperature growth. Having several versions of a gene could make differential regulation under different growth temperatures easier. For example, extra homologs (Kole_0111 and Kole_0110) of the earlier-discussed putative cold sensor system do not show coordinated temperature responses: Kole_0110 is up-regulated at 40°C, while Kole_0111 is up-regulated at 77°C (Table 1). Therefore, these additional homologs may encode sensors tuned to different temperatures.

### Evolutionary mechanisms that drive adaptive changes in Kosmotogales genomes

Gene family expansions can be achieved via within-lineage gene duplication or through LGT, and a combination of these mechanisms appears to be at work in *K. olearia*. For example, even though several Thermotogae genomes contain as many copies of PPIase genes as do *Kosmotoga* genomes (Table S9), phylogenetic analysis suggests that in the Kosmotogales this gene family has only recently been expanded by both LGT (the FKBP-type, Table 1) and duplication (the PpiC-type, Fig. S6C). Similar conclusions can be made from the phylogenetic analyses of *csp* genes (Fig. S6B).

LGT appears to be a significant factor in evolution of low temperature growth in Kosmotogales. In *K. olearia* although the proportion of transferred genes among the temperature responsive genes is similar to what is observed in the genome (Fig. S5), many of the genes identified here as key for low temperature growth have been acquired by LGT. For instance, the fatty acid synthesis genes with high expression at 40°C were acquired by LGT into the Kosmotogales and are likely important for low temperature growth in both *Mesotoga* and *Kosmotoga*. Other key LGT genes including the DEAD/DEAH box RNA helicase, the temperature sensor, and the low temperature induced *csp* genes, are only found in *Kosmotoga* spp. and may therefore be more important for its wide temperature growth (Table 1). The predicted acquisition of fatty acid synthesis and *csp* genes by (now mesophilic) archaea (López-García et al. 2015) additionally argues for the importance of these genes in adaptation to low temperature growth.

Despite the importance of expanded gene families in low temperature adaptation, the role of mutation and consequent natural selection on specific genes in response to changing environmental conditions should not be neglected. For example, typical cold response genes *rbfA* (Kole_2103) and *nusA* (Kole_1529) were not laterally acquired, but nevertheless show high expression only at 30°C. Deciphering adaptive changes that occurred in such genes compared to thermophilic homologs may elucidate molecular mechanisms of low temperature adaptation.

### Why maintain the capacity for growth over such a wide temperature range?

Most bacteria are under selection to eradicate extraneous DNA (and genes) from their genomes (Graur 2016), and among free-living bacteria Thermotogae in general have very compact genomes. Kosmotogales, however, have notably larger genomes than other thermophilic Thermotogae (Pollo et al. 2015;Zhaxybayeva et al. 2012), raising the possibility that expanded genomes are advantageous in *K. olearia*'s habitat. As discussed above, many of the genes in *K. olearia*, such as the cold-sensor system, were expressed only at specific sub- or supra-optimal temperatures, and do not seem to be important for growth at other temperatures (Table 1 and Table S5). The regulated response to low temperatures and the preservation of the laterally acquired genes specifically expressed at 40°C and 30°C suggest that *K. olearia* encounters environments with very different temperatures frequently enough to maintain these genes in its genome. Such environments may include oil reservoirs located at different depths, as well as marine sediments influenced by the mixing of cold deep-sea water and hydrothermal fluids (Sievert and Vetriani 2012). Perhaps, *K. olearia* migrates between such locations via subsurface fluids and, as a result, may have been selected to become a temperature generalist. Indeed, the environmental conditions of the subsurface environments and marine hydrothermal vents from which *Kosmotoga* spp. have been detected and/or isolated vary substantially (DiPippo et al. 2009;Nunoura et al. 2010;L'Haridon et al. 2014;Vigneron et al. 2017;Duncan et al. 2009;Duncan et al. 2014;Gittel et al. 2009;Kotlar et al. 2011;Pollo et al. 2016;Oko et al. 2017;Bordenave et al. 2013;Nesbø et al. 2010). Moreover, *Kosmotoga* sp. DU53, which is most similar to *K. olearia*, was isolated from an oil reservoir with an *in situ* temperature of ∼50°C, while *K. arenicorallina* was obtained from hydrothermal sediments with a temperature of ∼40°C (Nunoura et al. 2010). Notably, *K. olearia* was also identified as a major constituent of a metagenome from a deep subsurface oil reservoir with *in situ* temperature of 85°C and pressure of 25MPa (Kotlar et al. 2011). While the reservoir temperature is higher than the maximum *K. olearia* growth temperature reported here, elevated pressure could extend *K. olearia*'s temperature maximum, as has been demonstrated for some archaea (e.g. Takai et al. 2008). Therefore, *K. olearia*'s growth temperature range under natural conditions may be even broader than 20-79°C.

## Conclusions

The present study demonstrates that even bacteria with relatively small genomes can use transcriptional changes to respond effectively to large changes in temperature (Fig. 5). We showed that *K. olearia*'s response to sustained exposure to a non-optimal temperature includes up-regulation of hundreds of genes. Several key genes with known temperature-related functions apparently have been acquired laterally, suggesting that LGT is an evolutionarily successful strategy for expansion of temperature tolerance. However, gene duplication and subsequent neo-functionalization of the paralogs likely also play an important adaptive role.

The ability of *K. olearia* to inhabit both high and low temperature environments suggests that members of this lineage encounter environments with large temperature fluctuations and/or migrate across ecological niches within the deep biosphere (e.g. between deep and shallow subsurface oil reservoirs). Therefore, the subsurface environments, as well as their microbial populations, might be viewed as a connected archipelago instead of isolated islands. As a corollary, we speculate that *K. olearia-*like ecological generalists could also facilitate LGT among seemingly isolated deep biosphere microbial communities adapted to a narrower ecological niche. For example, we have previously demonstrated high levels of gene flow among hyperthermophilic *Thermotoga* populations in subsurface oil reservoirs and marine hydrothermal vents (Nesbø et al. 2015), environments that are separated by non-thermophilic surroundings but are hydrologically linked. The mechanism of such gene flow is not yet known, but *K. olearia*-like Thermotogae capable of growing both in subsurface oil reservoirs and adjacent marine sediments could serve as mediators of gene exchange.

Although some of the identified 573 temperature-responsive genes are already known to be expressed in bacteria and archaea grown at high or low temperatures, most of the up-regulated genes have not previously been implicated in temperature response and are in need of better functional and biochemical characterization. Moreover, the majority of the *K. olearia* genes responsive to elevated temperature encode proteins of unknown functions. Versatile proteins that work across a broad range of temperatures also warrant further biochemical and evolutionary analyses, as understanding of their enzymatic flexibility can aid the design of commercially important thermostable proteins.

## Material and Methods

### Bacterial culturing

*K. olearia* TBF 19.5.1 (DSM 21960(T), ATCC BAA-1733(T), Genbank accession number NC_012785) was grown at different temperatures (30°C, 40°C, 65°C, and 77°C), but otherwise optimal conditions, in *Kosmotoga* medium (KTM) using pyruvate as growth substrate as described in (DiPippo et al. 2009). Cultures used as inocula were stored at 4°C, except those used for experiments at ≥77°C. Actively growing cultures at temperatures ≥77°C had to be used directly as inoculum because the cultures would not grow from inocula stored at either 4°C or room temperature (∼22°C). Replicate cultures received the same volume of inoculum; however, variable inoculum volume was used at different temperatures (Table S1), as larger inoculum volumes were required to achieve growth at the non-optimal temperature treatments.

### Measurement of *K. olearia* growth at different temperatures

Growth curves were constructed from optical density measurements at 600 nm (OD_600_) using an Ultrospec 3100 pro. For cultures grown at 40°C, 65°C, 77°C, and 78°C two sets of triplicate bottles, inoculated from the same inoculum 12 h apart, were monitored for a 12 h period per day to generate the growth curves. The cultures for isothermic growth at 40°C, 65°C, 77°C, and 78°C were monitored hourly, while the cultures for shifted growth at these temperatures were monitored every 1-5 hours. At 30°C one set of triplicate bottles was monitored once daily. Isothermic growth curves were calculated from six replicates, except for the curves for 30°C and 77°C that had three and 12 replicates, respectively. All shifted growth curves consisted of six replicate cultures, except for 40°C and 30°C which had four and three replicates, respectively. To determine growth rates (Fig. 1), for each culture the lnOD_600_ was plotted against growth time and the curve was fitted with a linear trend line. The growth rate was defined as the slope at the log phase. To determine the time span of each growth phase (Fig. S1), full composite growth curves were constructed using pooled replicate data. For each curve, OD_600_ for all replicates was plotted against growth time and a polynomial regression trend line was fitted.

### Cultivation of *Kosmotoga* sp. DU53 and *K. arenicorallina,* and confirmation of their growth temperature ranges

*Kosmotoga* sp. DU53 was grown in KTM as described above for *K. olearia*. *K. arenicorallina* was also grown in KTM; however, maltose was used as substrate (2.5 mL and 0.5 mL 10% maltose was added to serum bottles and Hungate tubes, respectively). One mL of culture was used as inoculum for all cultures (bottles and tubes). The temperature range of each strain was assessed by examination of culture turbidity as a proxy for growth. Starting from cultures grown at optimal growth temperature (∼ 65°C for *Kosmotoga* sp. DU53 and 60°C for *K. arenicorallina*), new cultures were shifted in ≤10°C increments. If growth was observed after a shift, then that culture was used to initiate a new culture. The shifting procedure was terminated when growth was no longer evident at a given temperature.

### RNA isolation and processing

Cultures used for RNA extraction were inoculated from cultures that had been grown under the same temperature conditions for at least three transfers. The time at which a culture was expected to be in a desired growth phase was determined from the composite growth curves (Fig. S1), and was used as a cell harvesting time (listed in Table S1). This procedure avoided exposure of the cultures to the lower ambient temperatures in the laboratory during subsampling. For the 30°C cultures, OD_600_ was additionally measured 24 h before harvesting to ensure the culture was in mid log phase. In order to stabilize the transcripts and to avoid any transcriptional response to the change in temperature, an equal volume of “stop solution” (10% phenol in ethanol) was added to the sealed cultures via syringe immediately upon removal from the incubator. For each temperature treatment, RNA was extracted in mid-log to late-log phase, using the Zymo Research Fungal/Bacterial RNA MiniPrep Kit (Cedarlane Laboratories, Ltd.; Burlington, Ontario) and following the manufacturer’s protocols (Table S1).

Following recommendations in (Haas et al. 2012), we aimed to sequence ∼3 million non-ribosomal-RNA reads per sample. Ribosomal RNA (rRNA) depletion was performed on all samples using the Ribo-Zero rRNA Removal Kit (Gram-Positive Bacteria) Magnetic Kit (MRZGP126, Epicentre). On average, two aliquots containing 5ug total RNA (the maximum amount allowed by the Ribo-Zero kit) needed to be rRNA-depleted and then pooled to generate sufficient input RNA (200-500 ng for Ion Torrent PGM and 10-400 ng for Illumina MiSeq), although some samples required as many as five rRNA depletions. Quality and quantity of the total RNA, as well as efficiency of rRNA depletion, were assessed on an Agilent 2100 Bioanalyzer RNA Nano chip or RNA Pico chip following the manufacturer’s instructions for "Prokaryote Total RNA". RNA successfully depleted of rRNA was used to construct RNA-Seq libraries following the manufacturer’s instructions, and sequenced on either an Ion Torrent PGM (RNA-Seq kit V2; transcriptomes are labeled with an "IT" suffix) or an Illumina MiSeq (TruSeq RNASeq v2 2x100 bp). The platform and RNA extraction technique used for each transcriptome are summarized in Table S1. The transcriptomes are available in the Sequence Read Archive (http://www.ncbi.nlm.nih.gov/sra) under the accession number SRP075860.

### RNA-Seq analysis

For each transcriptome, sequenced reads were analyzed using the "RNA-Seq" function in CLC Genomics Workbench version 7.0.4 (https://www.qiagenbioinformatics.com/). Briefly, reads were first trimmed allowing no more than 2 ambiguous nucleotides and a maximum base-calling error probability of 0.05. To remove sequences that matched remaining rRNA transcripts, the trimmed reads were subjected to a relaxed mapping protocol: only rRNA genes were used as a “reference genome”, and reads were mapped only to the reference sequences if at least half of the alignment (length fraction = 0.5) had at least 50% identity to the reference (similarity fraction = 0.5); all other parameters were set to default (maximum number of hits for a read = 10, map to both strands, mismatch cost = 2, insertion cost = 3, deletion cost = 3, auto-detect paired distances. The mapped reads were designated as rRNA and were removed from further analysis.

The remaining reads were subjected to an RNA-Seq protocol with strict mapping parameters: at least 95% of the alignment had to have at least 95% identity (i.e. similarity fraction = 0.95; length fraction = 0.95) to the reference (the *K. olearia* TBF 19.5.1 annotated genome), mapping to intergenic regions was allowed, and all other default settings as described above. Unmapped reads were discarded. Expression levels for every gene were estimated using "<u>R</u>eads <u>P</u>er <u>K</u>ilobase of transcript per <u>M</u>illion mapped reads" (RPKM) values.

RPKM values for all genes are listed in Table S4. Differentially expressed genes were identified by doing pairwise comparisons of Illumina transcriptomes of the isothermically grown cultures at 30°C, 40°C, and 77°C to the cultures grown at the optimal temperature of 65°C. The analyses used the “Empirical Analysis of DGE” function in CLC Genomics Workbench, which employs the “Exact Test” for two-group comparisons (Robinson and Smyth 2008). A gene was considered differentially expressed in a pairwise comparison if it had (*i*) > 20 reads in at least one of the two transcriptomes, (*ii*) a statistically significant difference in the RPKM values (corrected for multiple testing using False Discovery Rate [FDR] < 0.05), and (*iii*) a difference in RPKM values at least two-fold in magnitude. Principal Component Analysis (PCA) and biplot visualization were performed using R packages *ade4* and *bpca* respectively (Dray et al. 2007;Faria et al. 2013). Each gene was assigned to a Clusters of Orthologous Groups (COG; (Galperin et al. 2015)) functional category using the Integrated Microbial Genomes (IMG) portal (Markowitz et al. 2014). Genes assigned to more than one COG category were counted in all assigned categories. Signal peptides and membrane domains were identified in Geneious v.9.

Batch culture cannot selectively discern gene expression that is exclusively influenced by temperature from expression that is solely growth rate-dependent. In theory, continuous culture conducted at a single growth rate could afford such discrimination. However, given the extremely slow growth of *K. olearia* near its temperature maxima and minima (Fig. 1), it was not feasible to use anaerobic bioreactors to cultivate cells at this marginal growth rate across the temperature range. Hence, in order to assess how differences in growth rates influence the expression patterns, we examined correlations of the expression pattern of the putative temperature responsive genes with growth rates calculated at each temperature. Pearson correlation of expression values and growth rates were calculated in Microsoft Excel. ANCOVA, linear regression, and likelihood ratio tests were carried out in R. The “growth rate only” model was the linear regression (expression = growth rate). In the ANCOVA model growth rates was set as the quantitative variable and temperature as qualitative variable (expression = growth rate + temperature). The growth rates used were 0.006 for 30°C, 0.087 for 40°C, 0.274 for 65°C and 0.107 for 77°C (see Fig. 1). When comparing the most significant temperature coefficient to the growth rate coefficient, the latter was scaled by the average growth rate. The growth rate effect was defined as being greater than the temperature rate effect if |temperature/(growth rate * 0.115)| < 1.

### Identification of genes involved in growth on pyruvate

*K. olearia* genes predicted to be involved in pathways for pyruvate conversion to acetate, CO_2_, H_2_ and ATP were retrieved from the KEGG (Kanehisa et al. 2014) and BioCyc (Caspi et al. 2014) databases. Genes encoding hydrogenases were taken from (Schut et al. 2012), and genes encoding the F-type ATPase subunits were identified using IMG (Markowitz et al. 2014).

### Fatty acids analysis

Total lipids were extracted from *K. olearia* grown at 40°C to early stationary phase and 65°C to mid-log, early stationary, late stationary and death phase by using methanol-chloroform (1:1 v/v). Fatty acid methyl esters (FAME) were prepared from total lipids extracts using mild alkaline methanolysis (Guckert et al. 1985). Dried FAME samples were re-dissolved in 300 μl chloroform (HPLC grade, Fisher Scientific) and analyzed by gas chromatography with mass spectrometry (GC-MS) on an Agilent 6890N gas chromatograph with a model 5973 inert mass selective detector (Agilent) fitted with an Agilent HP-5MS capillary column (30 m × 0.25 mm ID, 0.25 μm film thickness; J + W Scientific). Helium was used as the carrier gas with a temperature program of 130°C increasing at 3°C min^-1^ to 290°C and held for 2 min. Sample peaks were identified by comparison to Bacterial Acid Methyl Ester Mix standards (Supelco, Sigma Aldrich) or on the basis of mass spectra and expressed as a percentage of the total FAME quantified in each sample.

### Comparative analyses of three *Kosmotoga* spp. genomes

The genome of *K. olearia* was compared to genomes of *Kosmotoga* sp. DU53 (accession number JFHK00000000) and *K. arenicorallina* (accession number JGCK00000000) (Pollo et al. 2016) using the IMG portal (Markowitz et al. 2014) and Geneious v.9. Specifically, genes were declared homologous if they were significantly similar in BLASTP and TBLASTN (Altschul et al. 1997) searches (E-value < 10^−3^). For phylogenetic analyses, additional homologs of *K. olearia* genes were retrieved from the NCBI non-redundant (*nr*) protein database and the IMG databases via BLASTP searches, retaining the 100 top-scoring matches with E-value < 10^−3^. Sequences were aligned using MAFFT (Katoh et al. 2002), and phylogenetic trees were reconstructed using RAxML (Stamatakis 2006), as implemented in Geneious v. 9.1.3. Homologs from the recently released genome of *Kosmotoga pacifica* (NZ_CP011232) (L’Haridon et al. 2014) were included in gene-specific phylogenetic analyses. Pairwise Average Nucleotide Identity (ANI) (Goris et al. 2007) was calculated using the Enveomics Toolbox (Rodriguez-R and Konstantinidis 2016).

### Detection of laterally transferred genes

A customized version of HGTector (Zhu et al. 2014) (available through https://github.com/ecg-lab/hgtector) was used to identify putatively transferred protein-coding genes in the *K. olearia* genome. Homologs of each annotated protein-coding open reading frame (ORF) in the NC_012785 GenBank record were retrieved from a local copy of the NCBI *nr* database (downloaded November 21, 2014) using the BLASTP program from BLAST 2.2.28+ (Altschul et al. 1997). Sequences were first filtered for low complexity regions using the *seg* algorithm. Then, only matches with E-value <10^−5^ and sequence coverage ≥ 70% were retained. Database matches were expanded according to the *MultispeciesAutonomousProtein2taxname* file from RefSeq release 68. This was necessary as some genes across various taxonomic ranks were combined into a single entry in RefSeq, which artificially decreased the representation of these genes in Close and Distal groups (see below), and would confound downstream analysis. Taxonomic affiliation of each match was assigned using the NCBI Taxonomy database (downloaded on November 21, 2014). Only 500 top-scoring matches (after filtering for sequence coverage) were used as input for HGTector. The "Self" group was defined as TaxID 651456 (genus *Kosmotoga*), and the "Close" group was defined as either TaxID 1184396 (genus *Mesotoga*, a sister group) or TaxID 2419 (order Thermotogales, comprising *Thermotoga*, *Mesotoga*, and *Kosmotoga*). In either case, the “Distal” group comprised the remaining taxonomic groups. The conservative cutoff (the median between the zero peak and the first local minimum) was used for both the “Close” and “Distal” groups. A gene was designated as putatively transferred if its "Close" score was below the cutoff and its "Distal" score was above the cutoff. Putatively transferred genes with no top-scoring match in Thermotogae were designated as recent transfer events into *K. olearia* (labelled "K" in Table S8). Putatively transferred genes for which the difference between Close(Thermotoga) and Close(Mesotoga) scores was <1 were designated as gene transfer events into Kosmotogales (i.e. *Kosmotoga* and *Mesotoga*; labelled "K+M" in Table S8).

## Acknowledgments

This work was supported by a Research council of Norway award (project no. 180444/V40) to C.L.N.; a Simons Investigator award from the Simons Foundation, a Dartmouth College Walter and Constance Burke Research Initiation Award, and Dartmouth College start-up funds to O.Z.; and an NSERC Alexander Graham Bell Canada Graduate Scholarship CGS-M to S.M.J.P. We thank Rhianna Charchuk for technical assistance, Dr. Amanda J. Lohan for advice and discussion of RNA-Seq methodology, Dr. Karen Budwill for head-space gas analyses, and Jennifer Franks for initial discussions of lateral gene transfer detection methods.

